# Deep feature extraction of single-cell transcriptomes by generative adversarial network

**DOI:** 10.1101/2020.04.29.066464

**Authors:** Mojtaba Bahrami, Malosree Maitra, Corina Nagy, Gustavo Turecki, Hamid R. Rabiee, Yue Li

## Abstract

**Motivation:** Single-cell RNA-sequencing (scRNA-seq) has opened the opportunities to dissect the heterogeneous cellular composition and interrogate the cell-type-specific gene expression patterns across diverse conditions. However, batch effects such as laboratory conditions and individual-variability hinder their usage in cross-condition design.

**Results:** We present single-cell Generative Adversarial Network (scGAN). Our main contribution is to introduce an adversarial network to predict batch effects using the embeddings from the variational autoencoder network, which does not only need to maximize the Negative Binomial data likelihood of the raw scRNA-seq counts but also minimize the correlation between the latent embeddings and the batch effects. We demonstrate scGAN on three public scRNA-seq datasets and show that our method confers superior performance over the state-of-the-art methods in forming clusters of known cell types and identifying known psychiatric genes that are associated with major depressive disorder.

**Availability:** The code is available at https://github.com/li-lab-mcgill/singlecell-deepfeature

## 1 Introduction

Single-cell RNA sequencing (scRNA-seq) technologies profile the transcriptomes of individual cells rather than bulk samples [1, 2]. The wide adoption of scRNA-seq technologies enables the investigations of the molecular footprints at the unprecedentedly high-resolution for a wide spectrum of human diseases including cancer [3], autoimmune diseases [4, 5], Alzheimer’s disease [6], and major depressive disorder (MDD) [7]. However, single-cell data analysis still remains challenging due to confounding and nuisance factors, that manifest as individual variations or experimental biases such as different scRNA-seq technologies rather than biological variation. These confounding factors are often known as batch effects. Batch effects are the subsets of measurements that have different distributions because of being affected by laboratory conditions, reagent lots and personnel differences. [8]. The massive parallel sequencing [1] enable measurements with more than tens of thousands single-cell samples cross tens of human subjects in a single study (e.g., [3, 6]) further underscores the importance of addressing subject-level demographic confounders such as age and sex. Currently, there is a lack of highly scalable and robust model that enables systematic analysis of large-scale datasets while accounting for various confounding batch effects.

A number of methods have been developed for normalization, batch-effect correction, embedding, visualization and clustering of scRNA-seq gene expression profiles. [9] used mutual nearest neighbors (MNN) matching to account for batch effects. MNN operates on either the original space of the raw gene expression counts or the projected linear embedding space from the principal components analysis (PCA). However, MNN may be inadequate to model the non-linear effects known to exist in the scRNA-seq data [8]. Seurat [10] is another useful approach, which requires the users to carry out normalization, transformation, decomposition, embedding, and clustering of the gene expression samples in a step-by-step procedure. Each step is optimized empirically and independently of the other steps. Therefore, Seurat can be time-consuming to perform and it is a pipeline and not an end-to-end machine learning model. On the other hand, neural networks have demonstrated great promises in a wide range of domains and tasks such as computer vision, speech recognition, natural language processing, and computational biology [11]. Recently, neural networks have been exploited in scRNA-seq data mining. In particular, standard autoencoding neural networks were used to embed single-cell into low dimensional embedding without addressing the batch effect [12–14]. [15] described an autoencoder model for integrating cross-condition scRNA-seq data by learning their embedding via a random walk approach.

Variational autoencoders (VAE) [16] are efficient probabilistic models. Standard VAE assumes a Gaussian distribution both on the latent and observed variables, which is efficient to approximate the true posterior distribution by a variational distribution based on neural networks. VAEs are superior to standard autoencoders especially in modeling noisy data such as scRNA-seq profiles. This is attributable to their abilities to generate samples from the variational posterior distribution thereby approximating the marginal likelihood function in a principled Bayesian framework. However, in standard VAE, the generative distribution (i.e., encoder) *p*_*θ*_(*x*|*z*) is assumed to be Gaussian 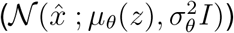, which is not ideal for modeling the scRNA-seq data. This is because the scRNA-seq experiment yield discrete integer counts with high dispersion, for which Negative Binomial (NB) are among superior and natural choices [17].

Recently, [18] and [19] used variational autoencoders as the generative frameworks for inferring the single-cell latent embeddings. A variational inference model called scVI proposed by [18] is the most relevant work to ours in terms of integrating both embedding and batch correction into one unified probabilistic model. However, scVI accounts for the batch effects by separating the batch variable from the rest of the biological variables in the latent space. Although this separation showed some improvements in practice, there is no guarantee that it will lead to a batch-free encoding of the single-cell samples without a more explicit constraint.

The new emerging paradigm of adversarial training has gained its momentum with the advent of Generative Adversarial Networks (GAN). GAN consists of a generative network and a discriminative network, both of which are optimized via a shared objective function corresponding to a minimax two-player game: the generator network tries to capture the true distribution by maximizing the loss function of discriminator [20]. Since its inception, many studies adopted the adversarial training procedure in different areas including visual domain adaptation [21] and single-cell modeling [22]. In particular, [22] utilized a GAN architecture to infer low dimensional single-cell embedding by jointly training a discriminator to distinguish fake scRNA-seq profiles generated from the generator from real scRNA-seq profiles. In contrast to our proposed scGAN model, their model does not correct for batch effects and is therefore unsuitable to model cross-condition heterogeneous scRNA-seq data.

In this paper, we present single-cell GAN (scGAN) to integrate both the single-cell embedding and batch correction procedure into a single unified and end-to-end deep generative model. One major contribution of our model is its unique ability to correct for continuous batch effects such as age, which is quite important in studying human subjects. We evaluate our model on several public scRNA-seq datasets including mouse/human pancreatic datasets and the recently available single-nucleus transcriptomic data from 17 MDD and 17 non-MDD subjects [7]. In all of these applications, scGAN demonstrates superior performance over the state-of-the-art (SOTA) methods in terms of clustering cells by known cell types, batch effect removal, and identifying biologically relevant and cell-type-specific genes that exhibit differential gene expression between MDD cases and healthy controls.

## 2 Methods

### 2.1 scGAN model details

#### 2.1.1 Model overview

Our proposed model consists of a Negative Binomial (NB) distributed variational autoencoder (NB-VAE) module to model the data likelihood of the scRNA-seq raw read counts and an adversarial-trained batch discriminator network to predict batch effect using the encoding from the NB-VAE module (**Fig.** 1). The objective of the NB-VAE network is to map the single-cell gene expression profiles into a low dimensional embedding, which dictates the sufficient statistics of NB data likelihood. To achieve this, the encoder needs to distill the most salient biological information from each scRNA-seq profile. Meanwhile, the encoder network also needs to minimize the correlation between its encoding and the confounding batch variable for each cell sample. The discriminator on the other hand learns to predict the batch variable using as input the embedding generated by the encoder network. The two network modules are learned in an adversarial fashion, each trying to improve themselves in adapting to the updates of the other opponent network. The following subsections contain the details of learning in each network module.

**Figure 1.**
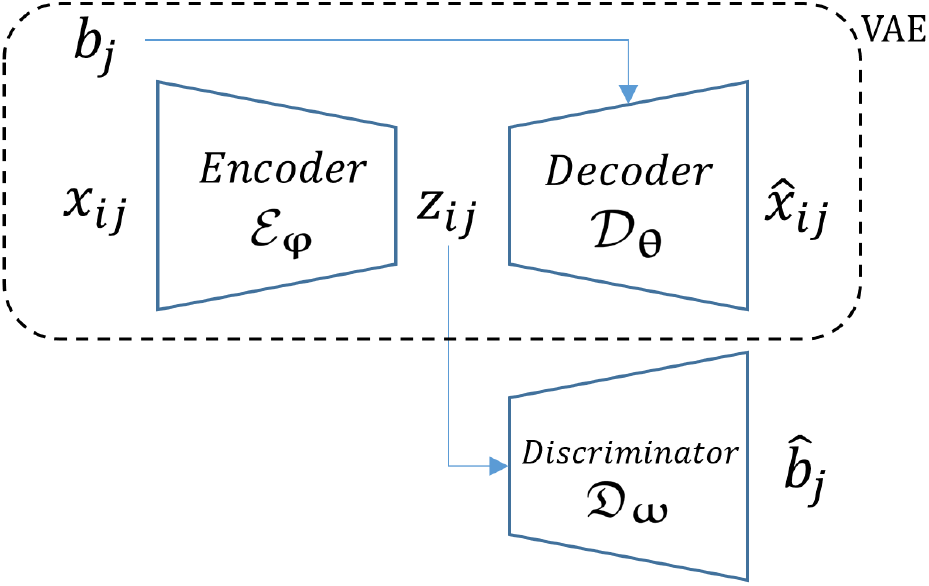
Single-cell Generative Adversarial Network (scGAN). The variational autoencoder (VAE) component of the scGAN model consists of the Encoder and Decoder networks. The Encoder projects each single-cell gene expression profile onto a low dimensional embedding. The Decoder takes the embedding as input and predicts the sufficient statistics of the Negative Binomial data likelihood of the scRNA-seq counts. The Discriminator, being trained adversarially alongside the Encoder network, predicts the batch effects using as input the Encoder’s embedding. Encoder, Decoder and the Discriminator are all parametric neural networks with learnable parameters denoted as **φ**, **θ** and **ω**, respectively. The input single-cell gene expression profile and the batch label for cell *i* in subject *j* are denoted as **x**_*ij*_ and *b*_*j*_, respectively. The reconstructed expression and the predicted batch label by the discriminator network for the input sample cell *i* in subject *j* are denoted as 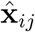 and 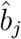, respectively. The latent variable as the gene expression embedding is denoted as **z**_*ij*_.

#### 2.1.2 Encoder

Each gene expression sample for cell *i* in subject *j* is an *N*-dimensional vector **x**_*ij*_ with the size equal to the total number of genes. We assume that the scRNA-seq data generative process is determined by a *K*-dimensional latent variable **z**_*ij*_, where *K ≪ N*. Our task is to infer the true posterior distribution of the latent variable *p*(**z**_*ij*_|**x**). We assume that the latent variable **z**_*ij*_ is sampled from a standard Gaussian distribution 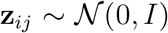. The marginal posterior 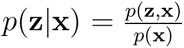 requires calculating the partition function by integrating out **z**: *p*(**x**) = ∫_z_ *p*(**x**|**z**)*p*(**z**)*d***z**, which is intractable. We approximate the true posterior *p*(**z**|**x**) with a variational distribution *q*(**z**|**x**), which is represented by the encoder neural network, i.e., 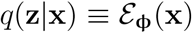. The marginal log-likelihood can be written as:

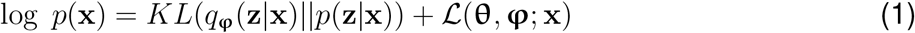

in which the first term is the KL divergence of the estimated posterior *q*(**z**|**x**) from real posterior *p*(**z**|**x**) and the second term is the evidence lower bound (ELBO):

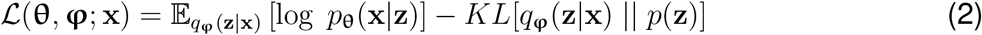

where the first term is the expected negative reconstruction loss and the second term is the KL divergence of the approximate posterior *q*_**φ**_(**z**|**x**) and the prior distribution *p*(**z**) of **z**. As the marginal likelihood log *p*(**x**) in Equation (1) is a constant, maximizing the ELBO 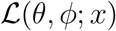 is equivalent to minimizing the KL divergence. The variational expectations in Equation (2) are approximated by taking the average over the randomly sampled data from the encoder network. The encoder network parameters are in turn optimized using stochastic gradient descent via back-propagation, which is made possible by the reparameterization trick due to the Gaussian latent variable [16]:

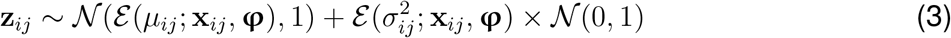

#### 2.1.3 Decoder

We use the decoder network 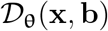 to model the sufficient statistics of a Negative Binomial (NB) distribution of the input data *p*(**x**|**z**, **b**), including the reparameterized mean and variance of the NB distribution:

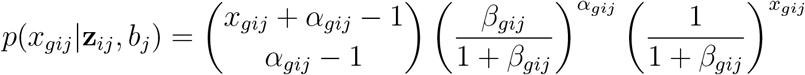

where *α*_*gij*_ and *β*_*gij*_ are the parameters for the NB function for cell *i* in subject *j* for gene *g*. We re-parameterize the NB function based on the mean *μ*_*gij*_ and variance 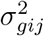 of the gene expression count, both of which are determined by the decoder neural network:

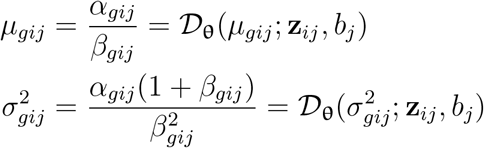

Notably, in order to generate realistic input gene expression 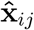, our decoder network uses both the embedding **z**_*ij*_ and the batch label *b*_*j*_ for cell *i* in subject *j* to model the NB parameters.

#### 2.1.4 Discriminator

We train a discriminator 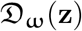 to predict the categorical or continuous batch effect label *b*_*j*_ based on the latent embedding **z**_*ij*_ for cell *i* in subject *j*, which are sampled from the 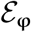 (Equation (3)). To create the adversarial interactions between the decoder and discriminator networks, we train the encoder to maximize the negative loss likelihood of the discriminator 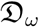. In other words, we optimize the parameters of the encoder and the discriminator in a `max min` function:

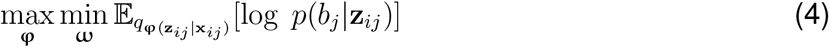

where *b*_*j*_ is the batch labels for single cell *i* that belong to subject *j* with scRNA profile measured as **x**_*ij*_. If the batch label is a discrete variable, we approximate objective function with the cross entropy (CE):

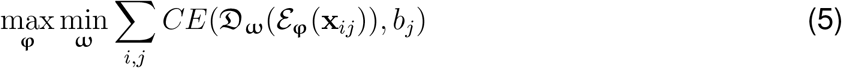

For continuous batch effect variable (e.g. the age of subject *j*), we use a univariate standard Gaussian to model its likelihood, which is equivalent to the negative sum of squared errors (*SSE*)

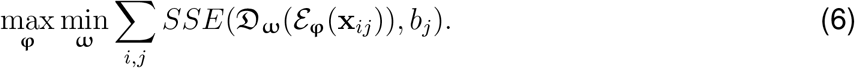

Optimizing the adversarial loss according to Equations (5) and (6) is proportional to simultaneously maximizing the ELBO while minimizing the correlation between the latent embedding and the batch label.

#### 2.1.5 Learning

To learn scGAN, we combine the ELBO in Equation (2) and the adversarial term of Equation (4) to give the following overall objective function:

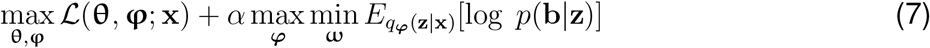

where 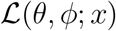 is the ELBO (2) and *α* is a free weighting coefficient that determines the importance of the adversarial term against the ELBO term from the VAE. We fine-tune the weighting coefficient *α* using a validation set.

In some applications, the parameters of the encoder network **φ** are optimized after training the batch discriminator network 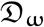. This means that the discriminator is trained until convergence given the encoder network. In our case, we do not wait until the discriminator is fully trained before updating the encoder in each iteration which leads to a faster training time.

#### 2.1.6 Deep differential analysis of single-cell gene expression

Due to small sample size in the scRNA-seq data, conventional hypothesis testing methods are often inadequate for identifying biologically meaningful differentially expressed genes (DEGs) that are associated with a biological condition of interest. For instance, our snRNA-seq MDD dataset only contain 34 subjects. To address this, we devised a powerful way by leveraging our deep neural network architecture to detect DEGs. For the ease of reference, we call this approach **scDeepDiff**. Our approach is inspired an early technique introduced in the deep learning community [23], which pre-trains an autoencoder on a large collection of unlabeled imaging data and then fine-tuning the pre-trained network weights for a classification task. Specifically, we first train our unsupervised scGAN until convergence. Then, we impose a two-layer feedforward neural network to predict whether a cell *i* comes from a subject *j* who has the disease phenotype (i.e., the 17 MDD cases among the 34 subjects in the MDD snRNA-seq dataset). In particular, the classifier uses as input the embeddings **z**_*ij*_ generated from the pretrained network by the encoder 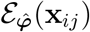 and outputs a probability from the logistic output unit as the prediction.

To assess the importance of each gene, we back-propagated the partial derivatives from the output unit of the network classifier all the way to the input units of the encoder, each representing an individual gene 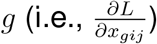. We aggregated these derivatives either within each cell cluster or over all cell clusters. These aggregated gradients indicate the importance of each gene in predicting the condition. We then sorted the genes by their aggregated derivatives and selected the top *M* genes as our predicted DEG candidates. This idea is inspired by DeepLift [24], which tackles a related but different problems from ours.

For scVI, we used their built-in function to perform the differential analysis. For Seurat, PCA, and MNN, we performed linear mixed model using the R package `lme4` [25] and `lmerTest` [26] to test DEGs within each of the clusters generated by these methods. In particular, we regressed the expression of each gene within each cluster on a set of covariates. We treated the biological conditions as fixed effects and the combined subject identifiers and batch identifiers as the random effects. The latter is to account for subject-dependent and batch-dependent variations that are distinct from the biological conditions. For each gene, we took either the sum or the maximum absolute test statistics (i.e., Bayes Factors for scVI and Wald-test statistic from LMM) across clusters to represent their association with the biological condition. We then chose the top *M* genes from the sorted gene list to represent the candidate DEGs.

### 2.2 Datasets

We evaluated our scGAN using three publicly available scRNA-Seq datasets with different types of batch effects as described below.

#### Mouse Pancreas Single-cell RNA-Seq Dataset

The mouse pancreas dataset contains 1,886 cells with 14,878 genes in 13 cell types after the exclusion of hybrid cell [27]. We downloaded this dataset from Gene Expression Omnibus (GEO accession number: GSE84133). The dataset was originally used to study the cell population structures of the human and mouse pancreas transcriptomic maps. The dataset contains populations from two distinct mice. We treated these two mice as two different batches because the distribution of their gene expression may be different due to their underlying conditions.

#### Human Pancreatic Islet Cells Dataset

The human pancreatic dataset consists of 6,321 pancreatic islet cells with 34,363 genes sequenced by four distinct sequencing technologies, CelSeq (GSE81076), CelSeq2 (GSE85241), Fluidigm C1 (GSE86469), and SMART-Seq2 (E-MTAB-5061) [28, 29]. The dataset was downloaded from the GEO database based on the above access numbers. Sequencing technologies have different cell dissociation and handling protocols. These technical variability can lead to strong batch effects. We used this dataset to demonstrate the capability of our scGAN in terms of multi-class batch effect correction.

#### Major depressive disorder single-nucleus RNA-seq Dataset

This is a newly available droplet-based single-nucleus transcriptomics dataset (GEO accession number: GSE144136). The dataset measures single-nucleus transcriptomes of 77,613 cells with 33,694 genes sequenced from the dorsolateral prefrontal cortex of 34 male post-mortum brains. Half of them are healthy controls and the other half are patients with Major Depressive Disorder (MDD) [7]. The number of cells is about the same for each subject. The subjects vary in ages ranging from 20 up to 80 years old. The age were distributed equally across cases and controls. We do not observe strong effects from the batch identifiers or subject identifiers in this dataset. However, the wide range of the subjects’ age prompted us to explore the benefits of accounting for age as a *continuous batch-effect variable* using our scGAN.

### 2.3 Experimental setting and scGAN implementation

For scGAN, we used a three layer neural network (two hidden layers and one output layer) for both Encoder and Decoder networks with 64 hidden units each. We set the latent space dimension to 10 for all the experiments. Our discriminator network was a two-layer network with 10 input units, which is the latent dimension of the embedding **z**, and a single hidden layer of size 20. We used Rectified Linear Unit (ReLU) [30] as the activation function for all of the hidden layers. The latent embeddings **z** are linear without the ReLU activation. The discriminator’s output is a softmax for discrete batch variables and linear function for continuous batch variables. We used mini-batch stochastic Adam [31] algorithm for optimization with the learning rate fixed at 0.001.

### 2.4 Evaluation

We used two criteria to measure the performance of our method in comparison with the SOTA methods. The first measure is *Batch Mixing Entropy (BE)*. BE measures how well the effect of confounding variable is removed from the latent embedding [18]. To calculate *BE*, we first randomly sampled a cell from the cell population. We then found the 50 nearest neighbors of the sampled cell based on the cell embeddings (which vary depending on the method). We measured the batch label frequency of the 50 nearest neighbor cells, which was used to calculate the entropy of the categorical batch variable. We repeated this for 100 randomly sampled cells and took the averaged results as the measurement for the BE.

By definition, if the distributions of *p*_***φ***_(**z**|**b**) are the same for each batch class label, we have a perfectly batch-corrected embeddings with the maximum BE. The range of BE measure depends on the number of batch labels (*k*_*b*_ = number of batches) and will be [0, log(*k*_*b*_)]. For continuous batch variables like the age of the subjects, we discretized the batch variable by dividing it into small 25% percentile bins and then measure the mixing entropy in the same way as for the discrete batch labels.

We also measured the clustering quality based on the *Adjusted Rand Index (ARI)*. ARI shows the similarity between the clustering found by our model using the cell embedding and the clusters of the ground truth biological cell-type groups. To calculate ARI, we needed to perform a discrete clustering on the cell embeddings derived from each method. We experimented with several clustering algorithms including k-means, Gaussian mixture modeling, and louvain cluster [32]. Our preliminary results showed that *louvain* clustering gave the most accurate clustering for all of the methods we compared, which is consistent to the published single-cell analysis by [33].

### 2.5 Method comparisons

To compare with scGAN, we chose three SOTA methods that can generate single-cell embeddings with batch-effect correction. In particular, we ran Mutual Nearest Neighbors (MNN) [9]), scVI [18], and Seurat [10] on the 3 benchmark datasets.

For the mouse pancreatic and human pancreas islet dataset, we provided the same batch identifiers to all of the methods including scGAN. Seurat, MNN, and scVI do not work with continuous batch effect variables such as age in the MDD dataset. Because the age range is between 18 and 87 years old, we grouped the subjects into 8 age groups from 10 years to 90 years with 10 years interval (i.e., 10-20, 20-30, …, 80-90). We then provided the 3 methods with the age groups as the batch class labels.

As a baseline method, we also ran simple PCA. To demonstrate the advantage and effectiveness of the discriminator network component of our approach, we also ran the same scGAN model but without correcting batch effects, i.e., removing the adversarial term from the objective function in Equation (7).

## 3 Results

### 3.1 Qualitative comparison of single-cell clustering

For the mouse pancreatic dataset, we first ran PCA followed by t-SNE visualization using the first 10 PCs. We observed that cells clearly cluster by the two animals rather than by cell types (**Supplementary Fig.** S1). Moreover, the human pancreatic cells projected by the first 10 PCs clustered by the scRNA-seq technologies rather than by the underlying cell types (**Supplementary Fig.** S2). To correct these batch effects in the data, we ran our scGAN along with the three SOTA methods (Section 2.5). The resulting clusters by each method exhibit improved separation of the known cell types (**Fig.** 2a and b, **Supplementary Fig.** S1 and **Supplementary Fig.** S2). Notably, none of the methods uses the ground truth cell types. They were evaluated entirely based on their unsupervised ability to cluster cells by their transcriptomes. By visual inspection, scGAN conferred relative larger separation of cell clusters by their cell types relative to the SOTA methods.

**Figure 2.**
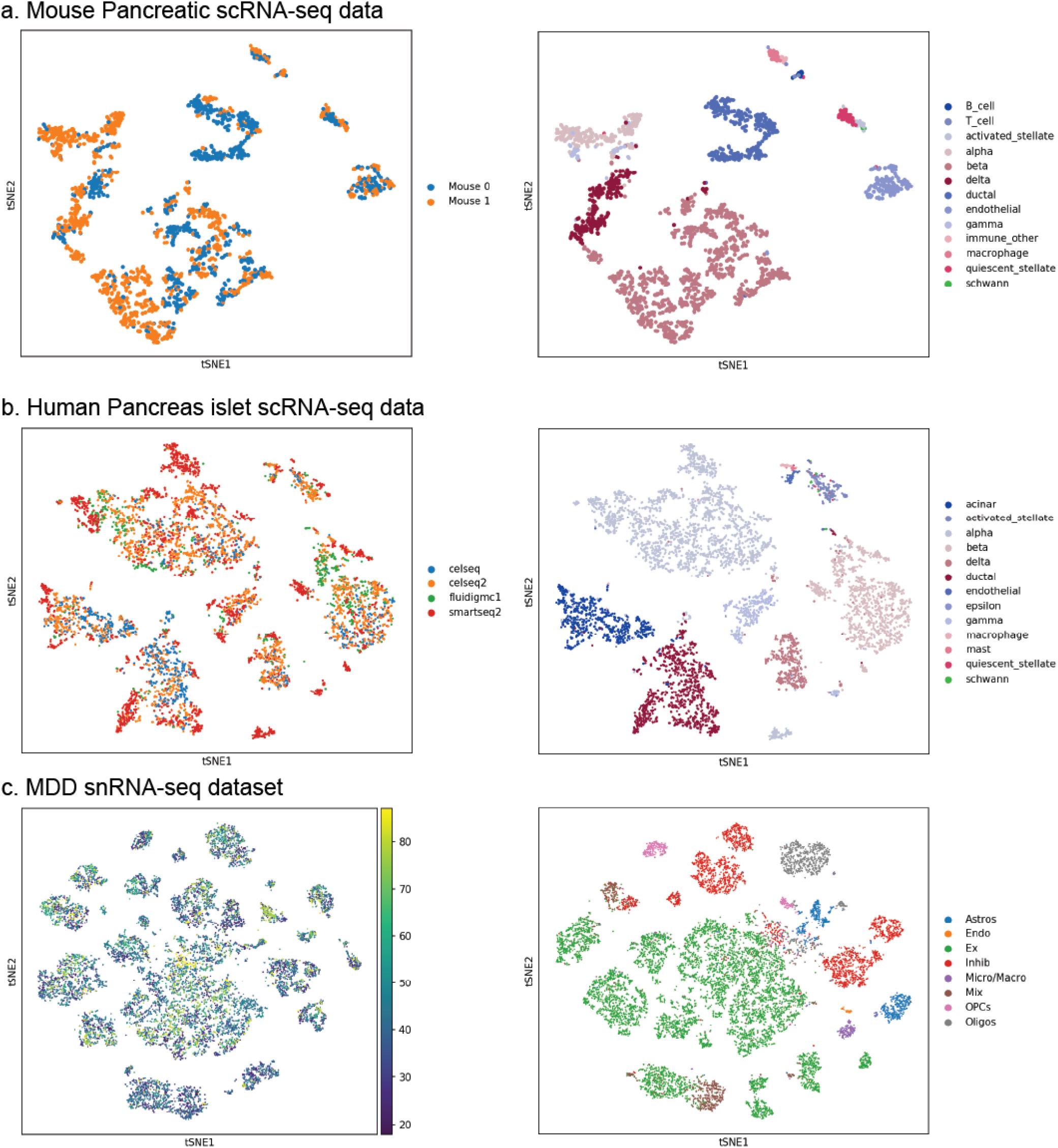
Cells clustering based on scGAN embeddings. **(a)** Mouse Pancreatic Cells dataset with the size of 1,886 cells from two mice, each treated as a batch. **(b)** Human Pancreatic Islet Cells dataset of size 6,321 cells sequenced by 4 different sequencing technologies of CelSeq, CelSeq2, Fluidigm C1 and SMART-Seq2. **(c)** MDD snRNA-seq dataset of size 77,613 confounded by the age variable of 34 patients. The left column shows the cells colored by batch labels. The right column shows the cells colored by the ground-truth cell types.

For the MDD dataset, although the distribution of ages are similar in cases and controls, we found that many cells tend to cluster by age when visualized based on PCA (Supplementary **Fig.** S3). This confounds our investigation on the cell-type-specific differential gene expression between the MDD case and control groups. Our scGAN accounts for the confounding age effects by the adversarial networks training. Here, we treated the age as a *continuous batch effect variable*. As a result, our scGAN discourages its encoder from generating embedding that can be used to predict age accurately by its discriminator, thereby generating cell embeddings of lesser age confounding effects (Section 2.1). Our approach led to clearer separation of cell types compared to the clusters generated by the baseline methods (**Fig.** 2c). Therefore, these qualitative results indicate the effectiveness of batch correction used by our method and also the higher flexibility of the non-linear embedding approaches compared with the linear PCA embeddings approach in general.

### 3.2 Quantitative comparison by clustering metric

We quantified the clustering quality based on Batch mixing entropy (BE) and Adjusted Rand Index (ARI) metrics (Section 2.4). scGAN compared quite competitively against all of the SOTA methods (**Fig.** 3). Also, we observed that the superior performance is largely attributable to the adversarial networks we introduced in our scGAN. In particular, when modeling the same dataset without using the adversarial component (scGAN^−^), we saw a drastic decrease in terms of both ARI and BE. In both mouse pancreatic and MDD datasets, our scGAN performs the best among all methods in terms of ARI. In human pancreatic islet data, scGAN is the second best led by Seurat. Notably, the human pancreatic islet was used in the original paper of Seurat [34], and it was possible that the method has been carefully fine-tuned on this dataset. We also observed that PCA has the worst performance in all 3 datasets, implying the linear approach is inadequate in modeling the scRNA-seq or snRNA-seq data. Surprisingly, the other deep learning model namely scVI (besides ours) does not perform very well on these three datasets. This is possibly due to the implicit way of modeling batch effects in scVI in contrast to our adversarial-network approach. Notably, scVI and our scGAN^−^ (without the batch correction) perform similarly in these three datasets. The overall clustering quality for mouse single-cell is higher than for the two human single-cell datasets possibly because of the homogeneous animals with identical genetic backgrounds and similar micro-environments compared to the human subjects.

**Figure 3.**
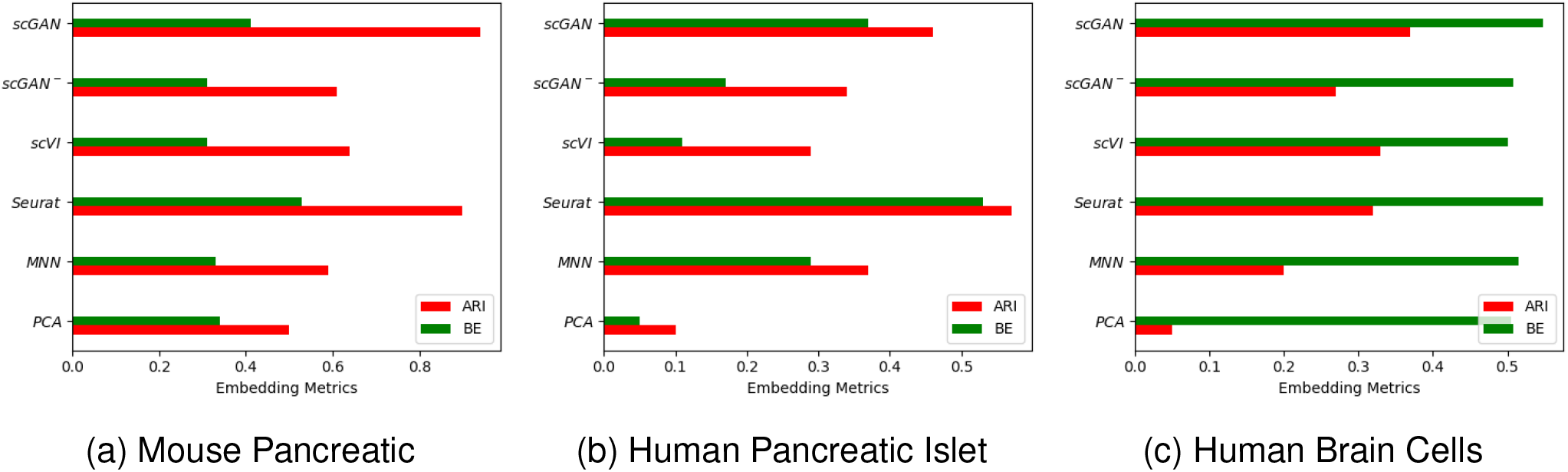
Comparison of clustering methods on 3 scRNA-seq or snRNA-seq datasets. Adjusted Rand Index (ARI) measures the consistency of embedding clusters with respect to the ground truth cell type groups. Batch Mixing Entropy (BE) measures the mixing of samples of different batches with each other. For both metric, the higher the better. The barplots display the ARI and BE scores for six embedding methods on the benchmark scRNA-seq datasets.

### 3.3 scGAN identified MDD-enriched cell clusters

We focused our analysis on the MDD snRNA-seq dataset because it is the largest snRNA-seq dataset by far on human psychiatric phenotype based on the Brodmann Area 9 (BA9) region of the patient postmortem brains [7]. We first sought to identify the cell clusters that were enriched for MDD subjects. Within each cluster, we computed the hypergeometric enrichment based on the number of cells that came from MDD patients relative to the total number of cells in that cluster and total number of MDD cells in the entire dataset. Out of the 10 clusters, we found two of them were significantly enriched for MDD (Hypergeometric p-value < 5e-25 and < 0.0086, respectively) (**Fig.** 4).

**Figure 4.**
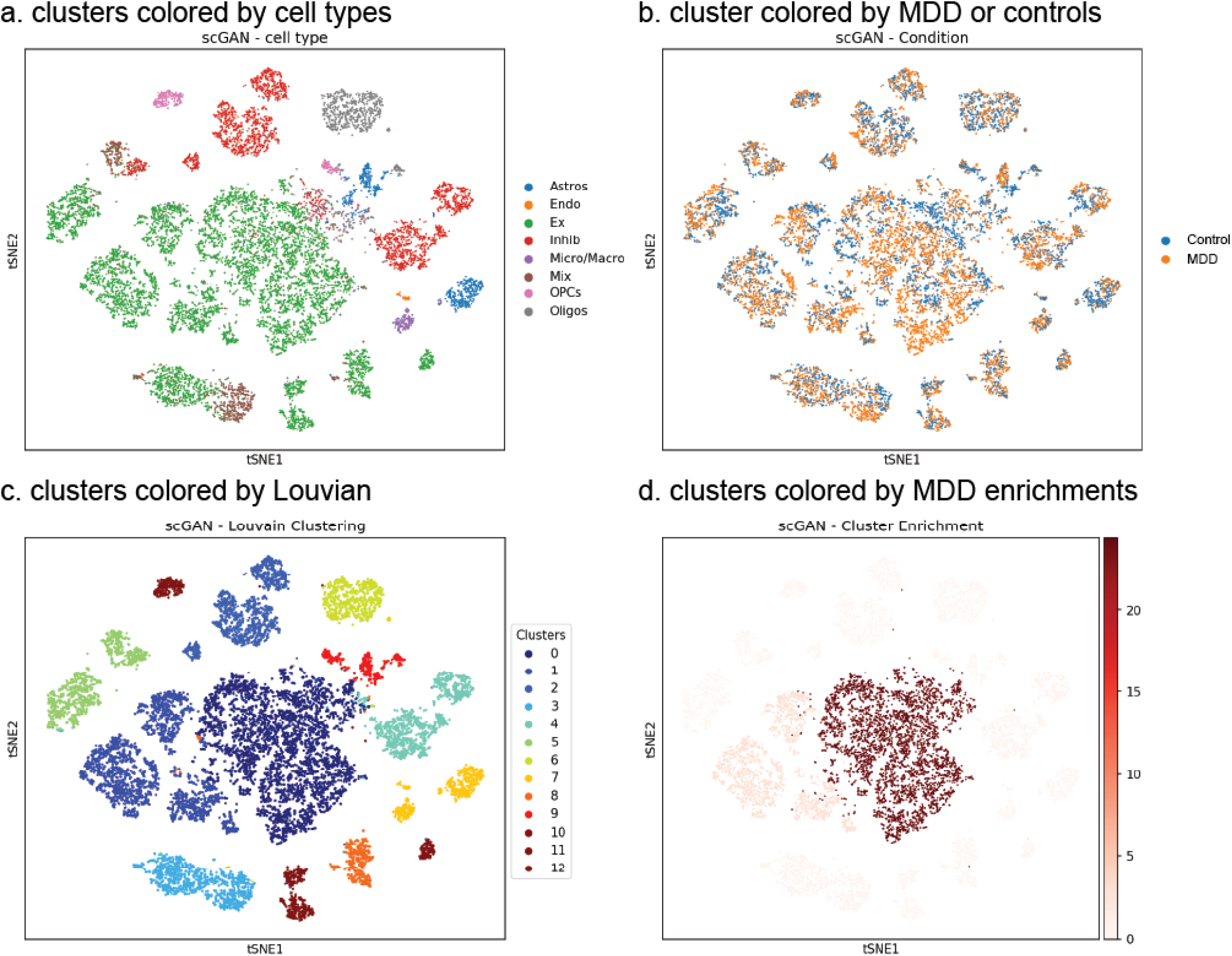
MDD-enrichment of cell clusters. We clustered the cells from MDD dataset and colored them based on **a**. cell types, **b**. MDD or controls, **c**. Louvain predicted cluster IDs, and **d**. MDD-enrichment scores. For each cluster, we tested whether the cells within the cluster are enriched for cells that came from the 17 MDD subjects. The statistical significance is assessed by hypergeometric test. The cells are colored by red with the intensity proportional to the -log_10_ p-values of the test.

Interestingly, both of these clusters (Cluster 0 and 1) are labeled with excitatory neuronal cell type (Ex). Indeed, this is consistent with the original findings by [7] via both empirical analysis and experimental validation. Besides ours, we found those clusters based on PCA, Seurat, and scVI each also yielded two MDD-enriched clusters (i.e., all of the methods except for MNN). All of these MDD-enriched clusters are also labelled as excitatory neuronal cell type. Therefore, these reassuring results demonstrated the consistency of our finding, in which the excitatory cells are significantly associated with the MDD phenotype among the 77,613 cells collected from the 34 male brains. However, because the excitatory cells represent the largest cluster, we must take caution in interpreting the biological implications of these results due to unbalanced number of cells per cell type cluster.

### 3.4 scGAN discovers more known MDD-associated genes

To identify differentially expressed genes (DEGs), we present a deep learning approach called scDeepDiff that uses the pre-trained weights by our scGAN to predict whether a given cell comes from one of the 17 MDD subjects or the 17 control subjects within each cluster. During the training, the gradients were back-propagated from the feedforward neural network to finetune the pre-trained weights for the prediction task. After training the model, we aggregated the error derivatives or gradients at each input gene of the classifer (i.e, the input units of the original encoder in the scGAN) (Section 2.1.6). For each gene, we further aggregated the gradients over all of the cells within each cluster or across all clusters to represent the importance of each gene in predicting MDD cels.

To validate our approach, we first took 90% of the cells for training and 10% for testing. For the largest cluster that were labelled as Excitatory (Ex) cell type, we achieved 78% accuracy on the testing cells, which is reasonably good provided the small sample size of only 34 subjects.

We then ranked the genes by the total sum of gradients across all clusters. We took the top *M* ∈ {5, 10, 50} genes and counted among these genes the number of PsyGeNET genes [35]. We found that there are more PsyGeNet genes found by our method compared to other approaches at every rank (**Fig.** 5; **Supplementary Table** S1). We also experimented using LMM on our scGAN clusters and found much fewer genes overlapping with the PsyGeNET genes: 1, 2, 7 PsyGeNET genes among the top 5, 10, 50 ranks. The results therefore suggest the improved power of using our scDeepDiff approach over the linear approach.

**Figure 5.**
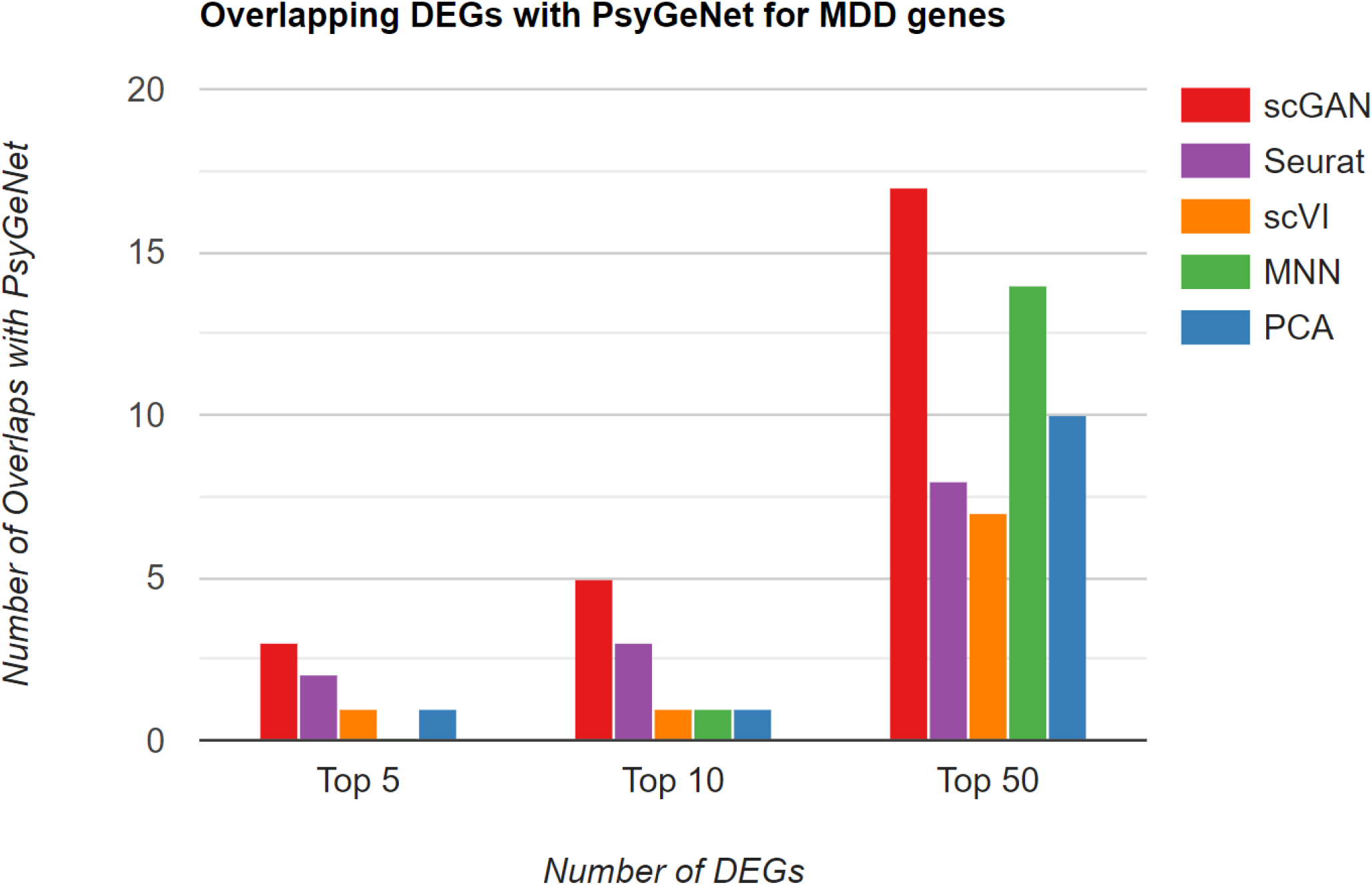
The number of PsyGeNet genes among the top ranked MDD-associated genes. We generated differentially expressed genes (DEGs) in the cell population collected from the 17 MDD subjects with respect to the cell population from the 17 controls. We then ranked the genes by the DE scores generated by each method. We chose the top *M* = {5, 10, 50} genes as the candidate MDD-associated genes and overlap them with the known psychiatric genes from PsyGeNet. The barplot displays the number of PsyGeNet genes found among the top rank genes by each method.

We further checked the PsyGeNet gene hits within each cluster. Here, we took top 50 genes *per cluster* and overlapped them with the PsyGeNet genes. Since only a few clusters are associated with MDD, we took the maximum number of PsyGeNet gene hits across all clusters as the representative score for each method. Once again, our approach ranks the best among all method with 22 PsyGeNet hits out of 50 top ranked genes, leading far in front of the second best method namely PCA and MNN (**Fig.** 6). Remarkably, the scGAN cluster that gives rise to the largest PsyGeNet hits is also the excitatory cluster that is the most significantly enriched for MDD among all of the clusters (**Fig.** 4).

**Figure 6.**
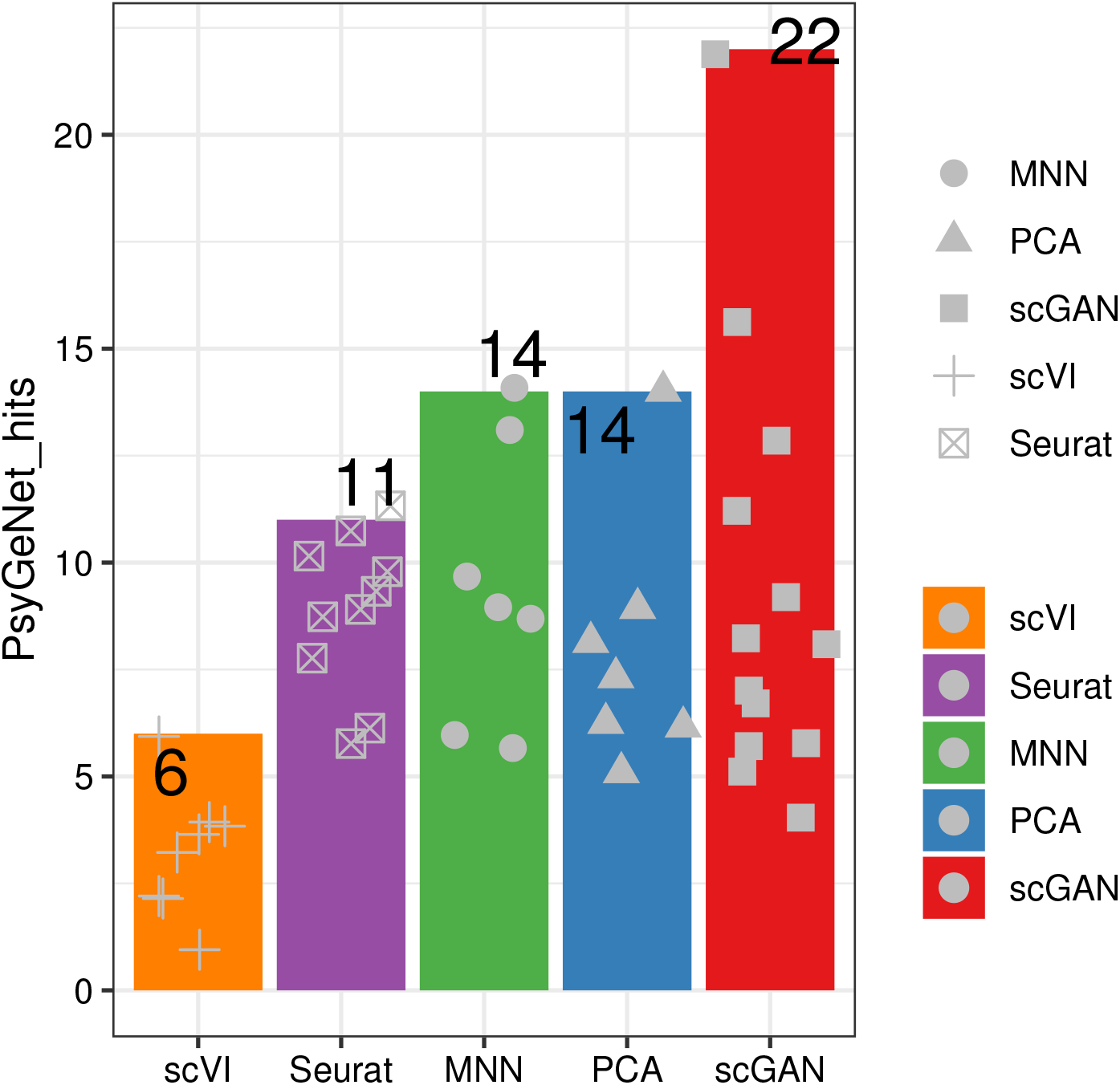
The maximum number PsyGeNet hits from one cluster. We generated differentially expressed genes (DEGs) in the cell population collected from the 17 MDD subjects with respect to the cell population from the 17 controls. We then ranked the genes by the DE scores. We perform this analysis in each cluster. We chose the top 50 genes per cluster and overlap them with the known psychiatric genes from PsyGeNet. The barplot illustrates the maximum number of PsyGeNet hits across all clusters for each method. The dots are the individual PsyGeNet hits for each cluster.

We also checked the overlap with reported genes that are associated with depressive orders in genome-wide association studies (GWAS). We obtained a list of 253 uniquely reported MDD GWAS loci from the GWAS catalog [36] with Experimental Ontology ID (EFO) EFO_0003761. We found 187 GWAS-associated genes that were also measured among the 33,694 genes in the snRNA-seq MDD dataset. Among the top 50 genes ranked by scDeepDiff, we found 3 genes namely *CLDN5*, *ITPKB*, *VCAN* that are associated with the MDD GWAS loci (hypergeometric enrichment p-value = 0.000173). Interestingly, the *ITPKB* loci is associated with Han Chinese ancestry MDD GWAS [37], whereas *CLDN5* and *VCAN* are associated with Scotland and/or UK Biobank MDD GWAS, respectively [38, 39]. In contrast, the top 50 genes from scVI, PCA, Seurat and MNN contain no MDD-associated GWAS genes. Overall, the overlap between transcriptome-wide DEGs and GWAS genes is small. Our results echos the low heritability of the MDD found in the recent GWAS [40] and underscores the importance of considering environments and epigenetic imprints that go beyond the germline mutations of the subjects.

## 4 Discussion

The advent of single-cell RNA sequencing (scRNA-seq) has unlocked the cell transcriptome at cellular-level resolution, providing theoretically optimal resolution potential. Extensive single cell surveys of tissues in mice [41] or humans [42] provide a portrait of variation in gene expression of cell-types in different tissue contexts. Compared to bulk RNA-seq, however, scRNA-seq is a much newer technology with more unknown technical biases, is much more expensive, is more challenging to perform, and is less sensitive to detect lowly expressed genes and prone to sequencing error [43, 44].

To address these challenges, we present scGAN as a deep embedding model to analyze scRNA-seq data. The main contribution of our method is the ability to simultaneously extract biologically meaningful embeddings and remove batch effects from each scRNA-seq profile. We demonstrated the utility of our approach on public scRNA-seq datasets. In particular, our approach does not only generate biologically meaningful single-cell clusters but also led to the discovery of MDD-associated cell types and MDD-associated genes. We attributed the success of our approach to the flexibility of the deep generative model and the adversarial network that we have introduced into the VAE framework.

As future works, we will further explore our approach in its ability to model massive scale single-cell sequencing data including cross-species, multi-omics (e.g., simultaneously modeling single-cell RNA-seq, single-cell ATAC-seq, and single-cell methylation while accounting for platform-dependent batch effects), multi-tissues, and multi-subjects. We will harness recently available human/mouse atlas data [41, 42] and disease-focused data such as scRNA-seq in patients with Alzheimer’s disease [6] and cancer patients tumors [45].

From methodological stand-point, like all of the neural network-based method, scGAN requires specification of the network architecture before training. In our preliminary experiments, we observed that scGAN is largely robust to the size of the architecture including the size and number of hidden layers in the encoder, decoder, and discriminator networks. This is attributable to the weight decay and dropout rates we have deployed in each network. Choosing the best network will require more extensive cross-validation ideally using a Bayesian optimization method [46]. Although our deep generative model demonstrates superior performance over some of the linear approaches such as Seurat and PCA, there is a further room for improvement in terms of its interpretability. In our future work, we will extend scGAN to incorporate prior biological annotations of cell-types-specific gene regulatory networks established from large reference atlas data such as the human scRNA landscape [42].

## 5 Conclusion

In summary, we demonstrate the utility of attenuating batch effects via an adversarial network while learning the low-dimensional single-cell embedding from the high dimensional scRNA-seq profiles. Our approach provides a crucial step towards discoveries of cell-type-specific differentially expressed genes in biological conditions of interest.

