## Supplementary Material for "Deep feature extraction of single-cell transcriptomes by generative adversarial network"

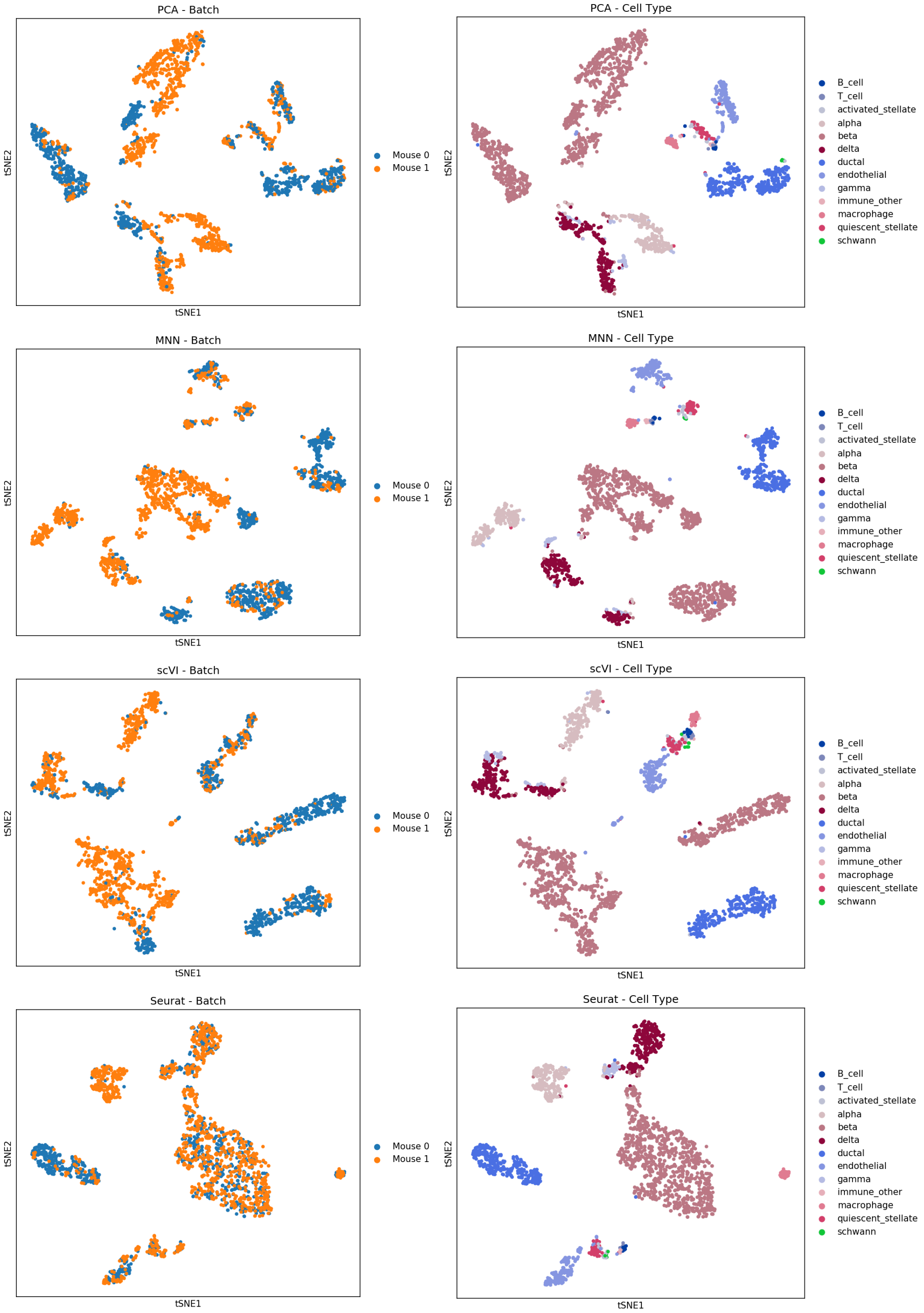


Figure S1 **Visualization of cell clusters on the mouse Pancreatic dataset using t-SNE based on the embeddings generated by each method.**

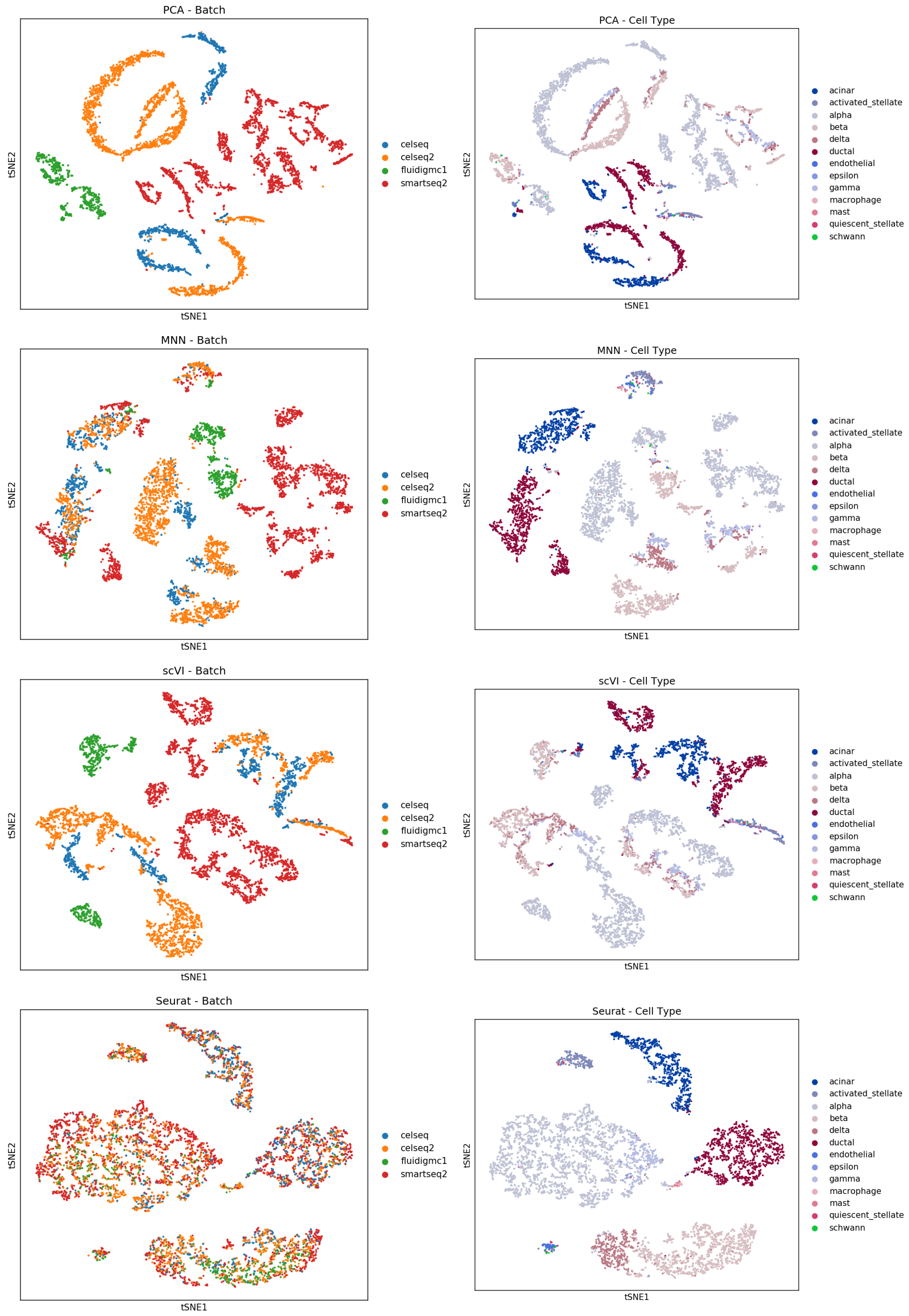


Figure S2 **Visualization of cell clusters on the human pancreatic dataset by t-SNE based on the embeddings from each method.**

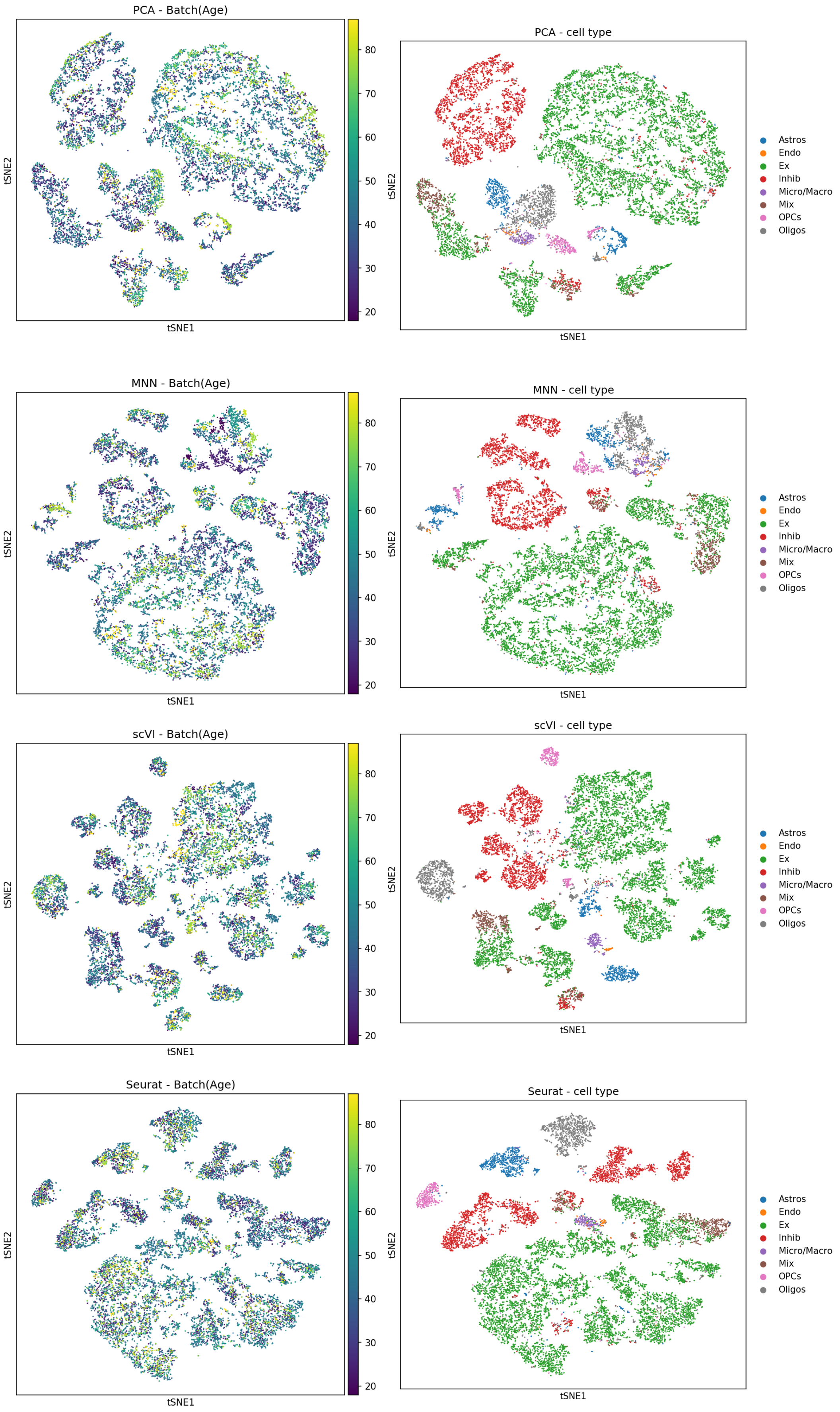


Figure S3 **Visualization of cell clusters on the snRNA-seq MDD dataset by the baseline methods by t-SNE on the embeddings from each method.**

Table S1: List of top 50 scGAN DEGs of human MDD brain cells dataset sorted by the the gradient score of each gene using our scDeepDiff approach.

| Gene | Gradient | Overlap with PsyGeNet |
|---|---|---|
| GLUL | 0.020525 | True |
| PTGDS | 0.020475 | True |
| FGFR3 | 0.019916 | |
| VCAN | 0.018657 | |
| EMX2 | 0.018364 | True |
| NDRG2 | 0.018054 | |
| LHFPL3 | 0.016145 | |
| KCNJ10 | 0.015293 | True |
| IGFN1 | 0.015266 | |
| APOE | 0.015164 | True |
| SDC4 | 0.015041 | |
| SLC27A4 | 0.014974 | |
| CTD-2017F17.1 | 0.014944 | |
| COL20A1 | 0.014868 | |
| IFITM3 | 0.014711 | |
| MLC1 | 0.014664 | True |
| LCAT | 0.014580 | |
| MT2A | 0.014541 | True |
| PCDH15 | 0.014453 | |
| SLC1A3 | 0.014203 | True |
| RP4-597A16.2 | 0.014057 | |
| NGFR | 0.013873 | True |
| GFAP | 0.013860 | True |
| LINC00609 | 0.013788 | |
| SLC1A2 | 0.013728 | True |
| DAAM2 | 0.013713 | True |
| MT1E | 0.013487 | |
| CARD14 | 0.013413 | |
| CH507-145C22.1 | 0.013327 | |
| ENTPD2 | 0.013287 | |
| TXNIP | 0.013256 | |
| CLDN5 | 0.013202 | True |
| RP4-668E10.4 | 0.013076 | |
| COL5A3 | 0.013074 | |
| RP1-253P7.4 | 0.013049 | |
| AP002856.7 | 0.013031 | |
| COL11A1 | 0.012865 | |
| GJA1 | 0.012855 | True |
| OLIG2 | 0.012535 | True |
| ATP1A2 | 0.012484 | |
| PLXNB1 | 0.012458 | |
| HAS2-AS1 | 0.012428 | |
| RP11-108L7.14 | 0.012420 | |
| C1orf56 | 0.012414 | |
| HLA-DQB2 | 0.012324 | |
| MSR1 | 0.012318 | |
| HEPH | 0.012251 | |
| SOX6 | 0.012200 | True |
| DBH | 0.012138 | True |
| CTD-2245E15.3 | 0.011978 | |

*Note*: Gene symbols, aggregated gradients scores over all clusters by our scDeepDiff approach, and whether gene overlap with PsyGeNet genes (True or empty) are shown in the table.
